# Contrasting carbon cycle along tropical forest aridity gradients in W Africa and Amazonia

**DOI:** 10.1101/2023.07.02.547401

**Authors:** Huanyuan Zhang-Zheng, Stephen Adu Bredu, Akwasi Duah-Gyamfi, Sam Moore, Shalom D. Addo-Danso, Forzia Ibrahim, Lucy Amissah, Riccardo Valentini, Gloria Djagbletey, Kelvin Anim-Adjei, Kennedy Owusu-Afriyie, Agne Gvozdevaite, Maria C. Ruiz-Jaen, Cécile A.J. Girardin, Sami Rifai, Cecilia Dahlsjö, Terhi Riutta, Xiongjie Deng, Minxue Tang, Yuheng Sun, Iain Colin Prentice, Imma Oliveras Menor, Yadvinder Malhi

## Abstract

Tropical forests cover large areas of equatorial Africa and play a significant role in the global carbon cycle. However, there has been a lack of in-situ measurements to understand the forests’ gross and net primary productivity (GPP and NPP) and their allocation. Here we present the first detailed field assessment of the carbon budget of multiple forest sites in Africa, by monitoring 14 one-hectare plots along an aridity gradient in Ghana. When compared with an equivalent aridity gradient in Amazonia using the same measurement protocol, the studied West African forests generally had higher GPP and NPP and lower carbon use efficiency (CUE). The West African aridity gradient consistently shows the highest NPP, CUE, GPP, and autotrophic respiration at a medium-aridity site, Bobiri. Notably, NPP and GPP of the site are the highest yet reported anywhere in the tropics using similar methods. Widely used data products (MODIS and FLUXCOM) substantially underestimate productivity when compared to *in situ* measurements, in Amazonia and especially in Africa. Our analysis suggests that the high productivity of the African forests is linked to their large GPP allocation to canopy and semi-deciduous characteristics, which may be a result of a seasonal climate coupled with high soil fertility.

## 3. Introduction

As the most productive terrestrial ecosystems, tropical forests and savannas account for over 60% of global terrestrial gross primary productivity (GPP) and feature large spatial variation ^1, 2^. Previous studies suggest that many African forests may have unique carbon budgets and dynamics, different to that of Amazonian forests^3^. This difference is likely because the two regions have experienced very different biogeographic and climatic histories and have different current environmental exposure ^3^. For instance, a satellite-based study suggested that the NPP trend of African forests during past decades has contrasted with that of Amazonian forests ^4^. Furthermore, a stable and positive carbon sink into above-ground biomass for African forests was reported to contrast with the declining trend in Amazonian forests, owing to the different mortality rates following drought events ^5^. A study focusing on the decadal long-term drying trend across West Africa also reported increases in forest biomass accompanied by a relative increase in deciduous species ^6^. Bennett et al. (2021) found that African forests were more resistant to drought than other tropical forests, likely linked to the long history of greater climatic variation in the African tropics, compared to Amazonia ^3^. Moreover, African tropical forests boast a higher biomass stock than neotropical forests ^8, 9^. Field measurements of NPP ^10^ revealed that along a wet-dry gradient in West Africa there was evidence that NPP was higher compared to equivalent gradients in Amazonia ^11^.

Despite its important and unique role in the global carbon cycle, our confidence in tropical primary productivity estimation, especially for African tropical forests, remains low ^12–15^. The large uncertainty is an inevitable consequence of the lack of African forest field evidence, which has been identified as a common issue in the carbon cycle community ^16–18^. For example, GPP can be estimated by non-biometric methods, such as eddy covariance tower measurements and remote sensing. However, in the entire African tropical forest region, there was only one eddy covariance tower (in Ankasa, Ghana), reporting three years of GPP (2011–2014) ^19, 20^. Satellite-based GPP products (for example MODIS-GPP) struggle to provide reliable estimates of GPP of the region owing to the dense cloud cover, which extends across both wet and dry seasons ^21–23^. Unlike GPP, autotrophic respiration (Ra) and allocation of GPP still require intensive field campaigns for quantification. To date, there has been no field assessment of GPP, Ra, GPP allocation and carbon use efficiency (CUE, the fraction of GPP allocated to NPP) of African forests. As most vegetation models calculate net primary production (NPP) and biomass from GPP and Ra ^24^, the dearth of these field measurements leads to simplified model assumptions or disagreement among models, restricting our ability to predict the effects of global change on the carbon dynamics of the biosphere ^25–28^.

To address these issues, we present the first comprehensive carbon budget for forest sites in Africa, along an aridity gradient and compare those to previously published results for an Amazonian aridity gradient ^11^. This dataset spans 14 one-hectare plots and six years of field data collection, including measurements of autotrophic and heterotrophic respiration that enable integrated assessment of GPP and its partitioning. Specifically, we ask: (1) is the higher NPP reported for the West African forests also reflected in higher GPP, or higher CUE? (2) Is the relationship between GPP and aridity similar along the West African and Amazonian forest gradients? (3) Do the studied West African and Amazonian forests share similar carbon flux partitioning patterns? (4) How do our in-situ measurements of tropical forest productivity compare with widely used global data products (MODIS and FLUXCOM – an eddy covariance tower based product)?

## 4. Comparison of productivity and respiration between Amazonia and West Africa

Our comparison reinforces that the carbon flux of the studied West African forests is distinctly different to that previously reported for Amazonian tropical forests. Overall, the total NPP and autotrophic respiration were higher in West Africa than Amazonia along the aridity gradients (Table S 2 and Figure 2), albeit the wettest sites share similar values. The difference arises mostly from differences in canopy NPP and leaf respiration (Figure 2). The high leaf respiration of West African forests is based on elevated dark respiration measurements, not on higher leaf area index (LAI) (Table S 1), consistent with previously reported high net assimilation rate and dark respiration of West African species ^29–32^. In contrast, stem woody productivity and rhizosphere respiration appear higher in the Amazonian studies, implying very different allocation patterns between the two studied regions (Figure 3). Overall, these differences lead to our estimates of GPP being much higher in the West African gradient than in Amazonia, particularly in the drier sites. However, unlike in-situ estimates, FLUXCOM (climate-based extrapolation from global flux tower networks) and MODIS (satellite remote sensing) estimated West African forests to have lower GPP than Amazonian ones (Figure 4). This is also evident in other studies using vegetation models or satellite-based products ^1, 33, 34^. We found FLUXCOM consistently underestimated both Amazonia and West African forests GPP (Figure 4), and MODIS GPP was even lower. It seems likely that dynamic vegetation models also underestimate West African forests GPP because previous studies involving our wet rainforest site Ankasa ^19, 20^ found model GPP estimates lower than the flux-tower GPP (yearly mean varying from 22 to 36 MgC/ha/year) that is smaller than in-situ biometric measurements (40.1 MgC/ha/year) ^35–37^. The above synthesis reveals an acute data-model discrepancy in tropical forests productivity estimates, especially for West African forests, which require more detailed investigation.

**Figure 1.**
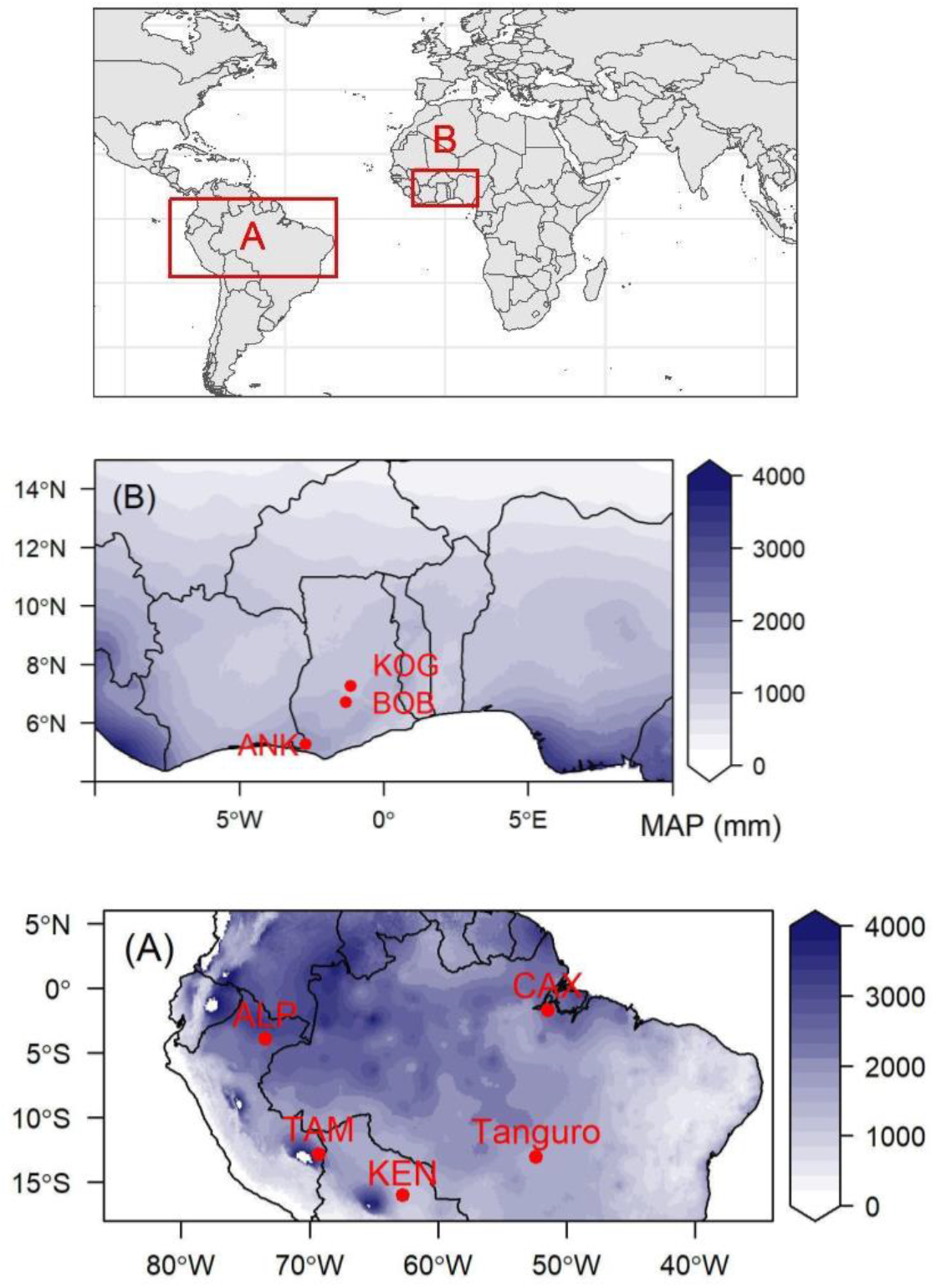
Map of study sites. (A) Amazonia aridity gradients and (B) West-African aridity gradient. Colour scale illustrates mean annual precipitation (MAP). Each red dot denotes a site. Each site contains multiple one-hectare plots (Table S1)

**Figure 2.**
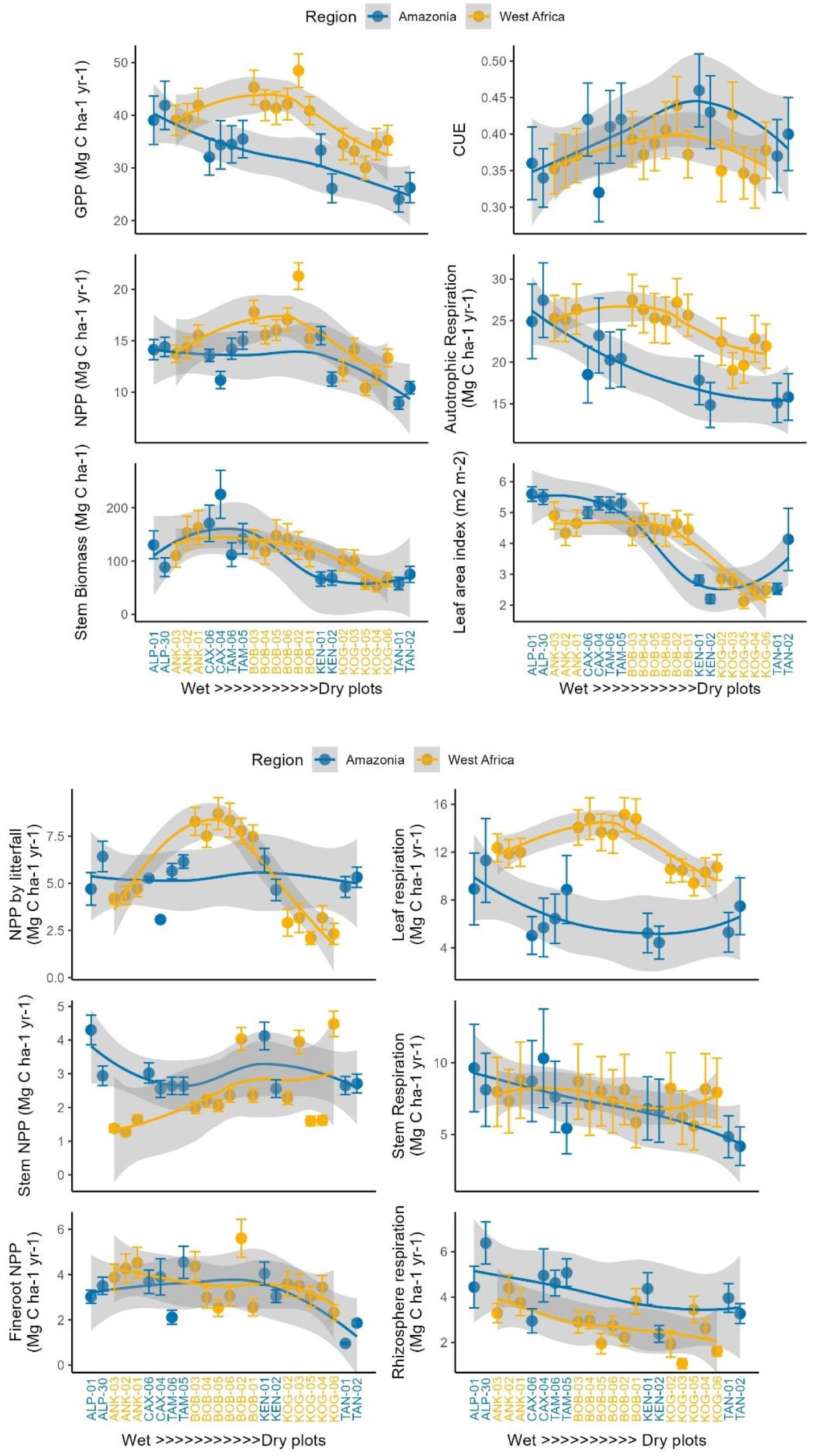
Biometric estimates of various components of carbon fluxes between Amazonia plots (red) and West African plots (green), including net primary productivity (NPP), gross primary productivity (GPP), and carbon use efficiency (CUE = NPP / GPP). See Table 1 for the definition of each carbon flux component. Local polynomial regression fitting lines (LOESS) are drawn for each region. Each dot represents one 1-hectare plot). The error bars represent the uncertainty of estimates (see Supplementary Material 1 for uncertainty estimation). The curves and their uncertainties (grey zone) were derived by Local polynomial regression. The x-axis is a factorial order of aridity. From wettest (left) to driest (right), sites were ranked by maximum climatological water deficit (MCWD mm year⁻¹). Within sites not discernible by MCWD, plots were ranked by in situ measured surface soil moisture (Table S1).

**Figure 3.**
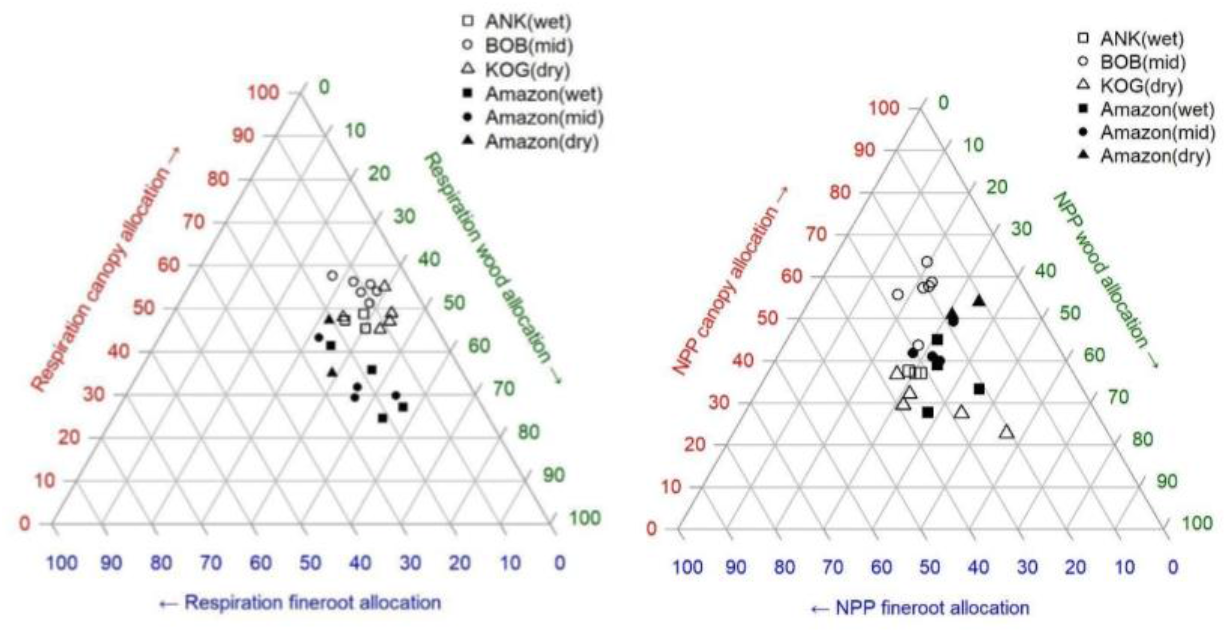
Percentage allocation of net primary productivity (NPP) and autotrophic respiration. One dot represents one plot, for West African (hollow marker) and Amazonia (solid marker). The aridity of plots was illustrated by marker shape. Allocation percentage should be read by following the arrow and ticks on each axis. For example, in the NPP diagram, the top hollow-round dot (BOB05) represents high allocation (63%) to canopy (red), 20% to wood (green), and 17% to fine roots (blue). The percentage of allocation was calculated as follows: NPP canopy allocation (%) * NPP = NPP_fine_litter_fall + NPP_herbivory; NPP wood allocation (%) * NPP = NPP_all_stem + NPP_coarseroot + NPP_branch; NPP fineroots allocation (%) * NPP = NPP_fineroot; Respiration canopy allocation (%) * R_autotrophic = R_leaf; Respiration wood allocation (%) * R_autotrophic = R_stem+R_coarse_root; Respiration fine roots allocation (%) * R_autotrophic = R_fine_root. See Methods for the definition of each component.

**Figure 4.**
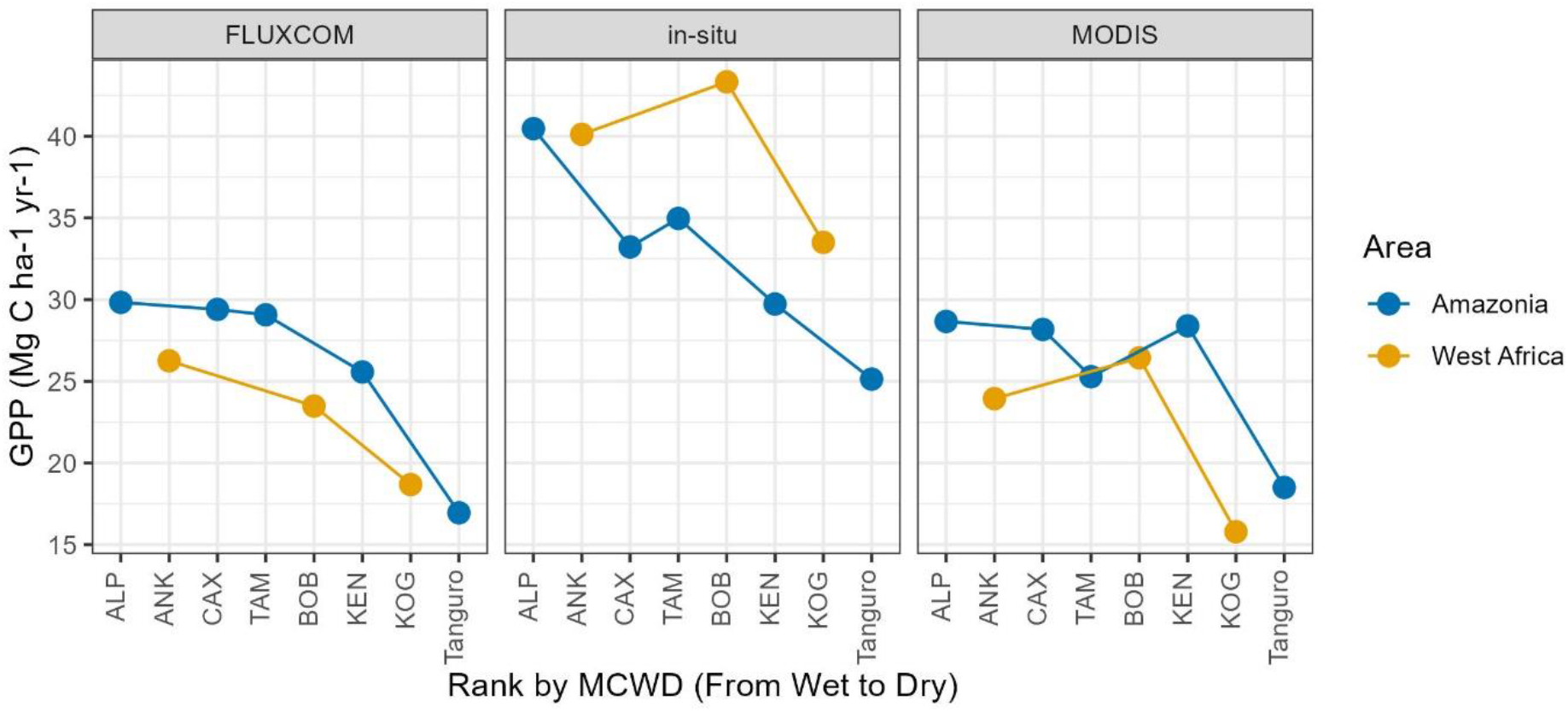
Comparing gross primary productivity (GPP, MgC ha-1 year-1) estimates of 5 sites in Amazonia (green dots) and 3 sites in West Africa (red dots) by FLUXCOM (left panel), in situ biometric methods (mid panel) and MODIS. The in situ panel is equivalent to Figure 2 but on a site scale. One dot represents one site; each site contains multiple one-hectare plots (Table S1)

## 5. Variation within sites and along the aridity gradients

It is striking to find that NPP is generally maintained, or even enhanced, in seasonally dry forests, before dropping off in very dry forests, along the aridity gradients in both Amazonia and West Africa (Figure 2). Along both transects, CUE peaks in the mid-aridity plots. This predominantly arises from patterns in leaf and fine root productivity. Woody stem productivity does show strong site-to-site variation, but no clear pattern along both aridity gradients. Surprisingly, along the African transect, GPP, autotrophic respiration, CUE and NPP are highest in the mid-aridity site, whereas GPP and autotrophic respiration decrease toward dry sites in Amazonia. In short, NPP follows the pattern of CUE and GPP in Africa but follows CUE not GPP in Amazonia. The pattern of GPP along the aridity gradient is well simulated in MODIS but not FLUXCOM (Figure 4). This study therefore highlights that dry forests can be more productive than wet evergreen forests, as revealed in the comparison between the West African mid-arid (BOB) and wet (ANK) forests, and the Amazonian lowland forests (NPP and GPP in Figure 2).

Within-site comparisons can reveal the impact of local environmental factors. For example, the seasonally flooded swamp forest ANK03, despite having less stem biomass (135.36 MgC/ha) than the adjacent dry substrate forest (162.7 and 153.3 MgC/ha for ANK01 and ANK02), does not have significantly different NPP and GPP (Figure S 4). Similarly, the different history of logging in Bobiri (see Methods), and steep variation of biomass along forest-savanna transition in Kogyae do not seem to result in any significant difference in either NPP or GPP (Figure S 4). There was also no discernible pattern when comparing photosynthate allocation between plots within any site (Figure 4). In contrast, previous studies in Amazonia reported that GPP and NPP change considerably from wet to dry sites ^11, 38, 39^ and along forest-savanna transition ^40^. A modest variation of NPP with logging intensity was reported in Borneo ^41^, but that was for much higher intensities of logging than at the Bobiri site. Such patterns were not seen for our Bobiri study and another West-African study on logging ^42^. The above implies that aridity is the primary driver of productivity and its partitioning of these forests. The lack of plot-to-plot variation increases the confidence of the exceptionally high GPP measurements found in Bobiri.

## 6. Contrasting patterns in carbon allocation

CUE in our African forest sites CUE appeared generally lower than previously reported values for Amazonian forests. Given that the Amazonia CUE is lower than the global average, our finding further expands the global range of CUE and confirms that, globally, mature tropical forests are low CUE ecosystems ^43–47^. Among all plots, CUE has no correlation with GPP (Figure S 1), whereas both GPP and CUE significantly correlate with NPP. Overall, CUE explains less spatial variation in NPP than GPP, but CUE does exhibit considerable spatial variation and assuming fixed CUE parameters in a model for both studied gradients would mimisrepresentpatial variation of NPP ^44^.

Further differences between Amazonian and West African forest carbon cycling were revealed in an investigation of photosynthate allocation into canopy, fine root, and wood NPP and respiration (see Figure 3 for definition). Note that this is the partitioning of productivity and metabolic activity instead of the more commonly reported partitioning of biomass ^48^. In both regions, the allocation pattern of autotrophic respiration is more homogeneous across plots than the allocation of NPP (i.e., points in Figure 3a are more clustered than in Figure 3b). For NPP, the Ankasa wet rainforest site shows allocation patterns within the spectrum of Amazonian plots. The mid-aridity site (Bobiri) consistently allocates more to the canopy than the reported Amazonian sites do. In contrast, the dry Kogyae site allocates consistently less to canopy and more to wood. For autotrophic respiration, leaf respiration clearly separates West African plots from Amazonian plots (Figure 2, 3). The dry Kogyae forests allocate more to wood respiration and the mid-aridity Bobiri sites allocate more to the canopy, both of which match the NPP allocation trends. Overall, the carbon allocation of tropical forests to different organs is highly variable between sites. Most variation can be seen in photosynthate allocated to the canopy, with a canopy respiration range from 24% to 63% and canopy NPP from 23% to 63%, substantially higher and more dynamic than that reported by previous extra-tropical meta-analyses ^49^.

The high variability in allocation patterns raises questions about the “fixed-ratio method” for intact forests carbon modelling ^50, 51^. Despite the large spatial variation, NPP partitioning to the canopy is the most dominant portion in our study site, consistent with previous in-situ evidence, but not shown in satellite-based products or vegetation models ^52, 53^. Moreover, along both the Amazonia and West-African aridity gradients, spatial patterns in woody NPP are poor proxies for the spatial pattern of total NPP or GPP – hence inferences on tropical forest productivity based on forest censuses alone should be treated with caution (Figure 2). Previous studies also found canopy NPP could explain greater spatial variation of total NPP compared with stem NPP ^54, 55^. The gathered evidence highlights the great importance of CUE and canopy NPP, whereas previous literature has paid much more attention to GPP and stem NPP ^56^; hence, future research is needed to balance and complete the carbon cycle.

## 7. Why are West African dry forests so productive?

As the wet evergreen forests in Amazonia Allpahuayo (ALP) and West African Ankasa (ANK) show similar levels of productivity (Figure 2), the reasons for the mean high productivity in the West African transect resides with the drier African plots (BOB and KOG). The most productive plot, BOB02 has a very high estimated GPP at 48.5 ± 3.2 and NPP at 21.3 ± 1.29 MgC/ha/year, the largest values reported this far for natural forests stands to our knowledge ^16^ (although, it should be noted that farmland or logged forest may have even higher NPP ^41, 42^). BOB02, compared to other plots in this study, has exceptionally high GPP, CUE, NPP, and autotrophic respiration simultaneously. All six plots in the Bobiri (BOB) Forest reserve (54 km^2^) show generally very high GPP (Figure S 2) so this high productivity is representative of the wider region. High NPP was also measured at another two one-hectare plots at Kakum, approximately 200 km to the south of Bobiri (BOB) ^42^ and by a previous study focusing on aboveground NPP ^57^.

Since the most distinctive difference between Bobiri forests and Amazonian forests is in carbon allocation to canopy (high leaf NPP and respiration), the high productivity of this mid-aridity site should be associated with its special semi-deciduous leaf phenology - remaining green all seasons but with only 5.2 ± 0.6 months leaf lifespan ^10^, substantially shorter than Amazonia (>12 months) ^58^. In other words, similar LAI and biomass but different turnover time, considering NPP = biomass x turnover. Such a carbon strategy, however, entails higher leaf-level photosynthesis to provide more carbon replacing leaves lost via turnover ^59^. Species at BOB and KOG were indeed characterized by high photosynthesis rates ^29, 30^. Along the West African aridity gradient, as indicated by photosynthetic traits measurements, photosynthesis rate per leaf area increases toward drier sites (which also have more sunshine) ^60, 61^ while LAI decreases (Figure 2), making mid-aridity site the most productive. In other words, light use efficiency increases toward drier sites, but fraction of absorbed photosynthetically active radiation (fAPAR) and shortwave radiation decrease. Similar to BOB02, another mid-arid plot in Amazonia, Kenia (KEN01), with the highest NPP along the Amazon aridity gradient, also featured a similar carbon strategy with more seasonality, higher CUE, and quicker carbon turnover. It is possible that BOB02 and KEN01 are recovering from previous disturbances (see Methods) ^39, 62, 63^ or that it is simply a common characteristic of semi-arid forests that experience higher rates of natural disturbance (drought or fire) and are dominated by fast-growing pioneer species ^64^. The high photosynthetic rate is consistent with short leaf lifespan and semi-deciduous phenology ^65, 66^. Nonetheless leaf traits provide the proximate factor explaining high productivity in seasonal African forests, not the underlying reason. This underlying reason is likely related to a combination of relatively high fertility (and associated cheap leaf construction costs) and seasonality favouring deciduousness.

In short, our findings suggest that semi-deciduous sites might be the most productive tropical forests, even more than wet evergreen. The counterintuitive finding is in fact consistent with ecological theories. In mature evergreen forests, each tree maximises its own resource acquisition leading to an increasingly conservative community that prioritizes defence and longevity over rapid growth”, leading to lower overall community resource use efficiency and lower primary productivity ^67–69^. In contrast, drier forests experiencing frequent dry spells are more subject to the intermediate disturbance hypothesis with higher mortality rates, species richness and productivity ^39, 70^. Previous studies reveal that long-term drying trends have driven these semi-arid forests toward more deciduousness ^6, 71^, perhaps making them more resilient to climate change than wet-evergreen Amazonia forests ^5, 7^. Our analysis highlights that West-African forests have a contrasting carbon strategy to Amazonian forests. However, as our sites are restricted to one country, more evidence is required to achieve a more geographically distributed understanding of the African forest carbon cycle.

In conclusion, we present the first complete carbon budget of African tropical forests to date by utilizing extensive field measurements across 14 one-ha plots, contributing towards a more comprehensive understanding of forest carbon allocation worldwide. The findings reveal that the productivity of West African mid-arid forests could exceed that of Amazonian lowland forests but was previously underestimated. Furthermore, the study features the discovery of the world’s most productive forests measured to date – Bobiri forest reserve in West Africa.

However, it should be noted that the Bobiri forest is now a mere 54 km^2^ patch. The West African tropical forests are severely fragmented ^72^ and increasingly face land use change pressures ^73, 74^. The forests not only present significant carbon stores ^8, 9^ and high productivity (Figure 2), but these forests are also potentially adaptable and resilient to change ^6, 75^. These irreplaceable and highly productive forests merit conservation and restoration attention to avoid continued carbon loss and for a multitude of benefits they provide to the local environment, to biodiversity and society.

## 9. Methods

### 1.1 The Ghanaian aridity gradient

As part of the Global Ecosystems Monitoring (GEM) network ^76^, 14 one-hectare plots were established within three forest reserves in Ghana (Table S1, Figure 1) spanning an aridity gradient. There are three plots in the wettest forest reserve, Ankasa National Park (ANK), which receives a mean annual precipitation of 2050 mm. The medium-aridity forest reserve Bobiri (BOB) hosts six plots with a rainfall of 1500 mm. At the driest end, Kogyae Wildlife Reserve (KOG) has five plots with a rainfall of 1200 mm. These plots have a very similar mean annual temperature, but span a steep precipitation gradient, which provided a “natural laboratory” to investigate the effects of aridity on forest productivity and respiration. The aridity of each plot is indicated by maximum climatological water deficit (MCWD mm/year) or, if not discernible by MCWD within a site, by *in situ* measured surface soil moisture (Table S1), in accordance with soil hydrology modelling of these sites ^61^). Along the aridity gradient, vegetation seasonality and deciduousness increased considerably toward the drier sites ^10^, whereas LAI and tree density decreased toward the drier sites ^61^. More information about the study sites, on soil properties, hydrology and climate regime can be found at ^10, 29, 77, 78^. Forests in ANK and BOB have never been burnt but KOG experiences wildfire roughly every decade ^19, 79^.

Although plots within the same forest reserve share very similar air temperature and precipitation, they differ dramatically in terms of soil moisture and composition of the vegetation community because of their soil properties, topography, and disturbance history. ANK is a humid rainforest and Pleistocene refuge with three plots spanning dry uplands (ANK01 and ANK02) and seasonally inundated riverine lowland (ANK03). BOB is a semi-deciduous forest where the plots span a gradient with selective logging history, ranging from the intact forest (BOB01) to forest lightly logged (2-3 stems/ha extracted) in 1959 (BOB02 – 04) and in 2001 (BOB05 -06). KOG is a forest-mesic savanna transition where local soil factors appear to influence vegetation type from dry forest (KOG02 and 03) to savanna (KOG05and 06) ^61^. Within any site (e.g., within ANK), there are common species between plot but the most abundant species may still differ; there are almost no common species between sites.

### 1.2 Detailed carbon budget quantification

The study quantified the whole carbon cycle of 14 West African plots (one-hectare each) with a bottom-up method mostly following the GEM protocol ^76, 80^. For each GPP component, a definition and brief description of the sampling method is provided in Table 1. The detailed sampling technique, calculation and scaling process, and references are explained in Supplementary Method. For completeness and consistency, we sourced the estimate of some NPP components from a previous analysis of the same dataset ^10^. Carbon allocation (the partitioning of GPP) was illustrated with a ternary plot (Figure 3), drawn by R package “Ternary.” The Z test was used to assess the difference among plots (Figure S 4).

**Table 1.** Abbreviation for components of NPP. See Supplementary Method for field protocol and calculation procedures. Data were retrieved from ^10^ except NPP_root_exudate.

| Components of NPP | Meaning | Sampling period and interval |
| --- | --- | --- |
| NPP_canopy | The sum of NPP_fine_litter_fall and NPP_herbivory |  |
| NPP_fine_litter_fall | NPP by litterfall in Figure 2, measured by litter fall trap, is the sum of leaf, twig, flower, fruit, seed and 'unidentified', not including NPP_herbivory and NPP_branch. The sum of flower, fruit and seed is called 'reproductive NPP'. | 2011–2013(ANK)<br>2012–2015(BOB&KOG),<br>every 14 days |
| NPP_herbivory | NPP_leaf multiplied by 'the percentage of leaf bitten by animals', which were obtained from leaf scan. | 2012–2013(ANK)<br>2012–2015(BOB&KOG),<br>every two months |
| NPP_branch | Measured with coarse woody debris census (i.e. branch litter larger than fine litter fall), it is representing the NPP that trees used to update their branch every few months. The branch is growing larger as the tree stature enlarge, meanwhile, the small branch is also updating themselves (e.g. because of wind damage), the former one is counted in NPP_all_stem, the later one is counted in NPP_branch | 2012–2013(ANK)<br>2012–2015(BOB&KOG),<br>every three months |
| NPP_big_stem | Measured through stem diameter and tree height, for tree larger than 10cm diameter at breast height, representing the expanding of tree stem | 2011–2013(ANK)<br>2011–2015(BOB&KOG),<br>every year |
| NPP_small_stem | Measured through stem diameter and tree height, for tree smaller than 10cm diameter at breast height, representing the expanding of tree stem | 2011–2013(ANK)<br>2011–2015(BOB&KOG),<br>every year |
| NPP_all_stem | The sum of the above two. This is the 'Stem NPP' in the main figure |  |
| NPP_coarseroot | Estimated by a root shoot ratio of 0.21 multiplied by NPP_all_stem, representing the expanding of tree coarse root | Estimated |
| NPP_fineroot | Measured using 'ingrowth core' technique, representing NPP in reproducing fine roots | 2012–2013(ANK)<br>2012–2015(BOB&KOG),<br>every three months |
| NPP_herbs | Measured by grass harvest, NPP of understory vegetation, mostly herb and grass | 2014–2015(KOG<br>only, there are little herbs<br>in BOB and ANK), every<br>three months |
| NPP_root_exudate | NPP of root exudates, estimated as a mean of several methods <sup>84</sup>. See Supplementary Method. | Estimated |
| NPP_AG | Aboveground NPP, sum of NPP_fine_litter_fall, NPP_herbivory, NPP_branch, NPP_all_stem and NPP_herbs |  |
| NPP_BG | Belowground NPP, sum of NPP_fineroot, NPP_coarseroot and NPP_root_exudate |  |
| NPP | The sum of NPP_AG and NPP_BG |  |
| NPP_voc | NPP of volatile organic compounds, not included in this study. It is thought to be a negligibly small proportion of total NPP <sup>54</sup>. | Not included |

**Table 2.** Abbreviation for components of respiration and detritus flux. See Supplementary method for field protocol and calculation procedures.

| Components | Meaning | Sampling period and interval |
| --- | --- | --- |
| R_leaf | Respiration flux from leaf surface, calculated as $R_{\text{dark}} \times \text{LAI}$ | |
| LAI | Leaf area index, measured with hemispherical photo | 2016-2017(ANK), 2012-2017(BOB&KOG), every month |
| Rdark_shade | Measured leaf dark respiration for shade leaves | 2014-2016 (ANK, BOB & KOG), every three months |
| Rdark_sun | Measured leaf dark respiration for sun leaves | 2014-2016 (ANK, BOB & KOG), every three months |
| R_stem | Measured stem respiration | 2016 (ANK), 2014-2016 (BOB & KOG), every month |
| R_soil | Soil respiration, including litter, but not coarse root respiration. The components were measured with EGM5 soil chamber, We assume that the soil respiration measurement (by installing a chamber on the ground) will capture fine litter respiration, soil organic matter respiration, rhizosphere respiration (including fine root respiration, mycorrhizal and priming effect etc), but will not capture coarse root respiration<sup>82</sup>. We then break this down into R_fine_litter, R_SOM and R_rhizosphere with partitioning experiments (Riutta et al., 2021). | 2016 (ANK), 2013-2016 (BOB), 2014-2016 (KOG), every month |
| R_rhizosphere | Respiration associated with rhizosphere (like fine root, mycorrhizal or priming effect), see R_soil for method | Derived from the above with partitioning experiments |
| R_fine_litter | Respiration from litter layer above soil, see R_soil for method | Derived from the above with partitioning experiments |
| R_SOM | Respiration from soil organic matter, see R_soil for method | Derived from the above with partitioning experiments |
| R_soil_heterotrophic | The sum of R_fine_litter and R_SOM | Derived from the above with partitioning experiments |
| R_coarse_root | Coarse root respiration, estimated as root shoot ratio multiplied by R_stem | Estimated |
| R_autotrophic | Total autotrophic respiration = $R_{\text{stem}} + R_{\text{coarse\_root}} + R_{\text{fine\_root}} + R_{\text{leaf}}$ | |
| R_heterotrophic | Heterotrophic respiration, The sum of R_cwd and R_soil_heterotrophic, by definition should also include animal respiration and R_mycorrhizal but these were not included in this study. |  |
| R_cwd | Respiration from coarse woody debris (any necromass not included as fine litter), estimated in this study | Estimated |
| GPP | Gross primary production, as $NPP + R_{autotrophic}$ | |
| CUE | Carbon use efficiency, as $NPP/GPP$ | |
| NEP | Net ecosystem productivity (Same as net ecosystem exchange), as $NPP - R_{heterotrophic}$ | |
| D_litter_fall | Detritus in the form of fine litter, from aboveground biomass to soil, equal to $NPP_{fine\_litter}$ | |
| D_CWD | Detritus in the form of coarse woody debris, from aboveground biomass to necromass, as $NPP_{all\_stem} + NPP_{cwd}$ | |
| D_root | Detritus from both coarse roots and fine roots to soil, as $NPP_{fineroot} + NPP_{coarseroot}$ | |
| D_CWD_to_soil | Detritus from CWD to soil |  |

Our reported NPP and R_autotrophic are completely independent measurements. The plot-to-plot variations (Figure 2) in NPP and R_autotrophic are highly similar, and analogous allocation patterns (Figure 3) could also be found between NPP and R_autotrophic, which reinforce the reliability of the findings. Furthermore, we found the sum of R_soil_heterotrophic and R_cwd was roughly equal to the sum of D_cwd, D_litter_fall, and D_root in each site (Figure S 3). This is a valuable cross-check to validate our carbon flux measurements because R and D are independent measurements, and they are expected to be equal in steady-state conditions.

The study also featured a comparison with detailed carbon budgets of Amazonian plots where the same sampling protocol was applied ^76^. The carbon budget of these plots is reported by ^11^. The western Amazonian sites (on relatively fertile soils) include Allpahuayo in NE Peru with no seasonality ^81^, Tambopata in SE Peru with a moderate dry season ^82^ and Kenia in Bolivia featuring strong dry season ^39^ which is situated at the transition between humid Amazon forest and *chiquitano* dry forest. The eastern Amazonian sites (on relatively infertile soils) include Caxiuanã, humid forests in NE Brazil ^83^ and Tanguro dry forests in SE Brazil ^64^, which sit close to the dry forest-savanna ecotone. See Table S 1 and references above for more characteristics of the sites.

### 1.3 FLUXCOM and MODIS

Using Google Earth Engine, we retrieved MODIS GPP from the collection MOD17A3HGF during 2001 to 2020. This collection of GPP has been cloud contamination filtered and gap filled by the data providers. We chose the ‘RS_METEO’ version of FLUXCOM because the magnitude of GPP in this version does not involve uncertainty from MODIS FAPAR, which makes the comparison between FLUXCOM and MODIS GPP more independent. We extracted GPP of the studied plots using their coordinates and calculated mean per site.

## 2 Acknowledgement

We thank Sophie Fauset, Roberto Salguero-Gómez, Simon Lewis, Jesus Aguirre Gutierrez, and David Bauman for valuable suggestions. We thank Armel T. Mbou, Agnes de Grandcourt and Giacomo Nicolini for their help with field works. Y.M. is supported by the Jackson Foundation. H.Z. received Henfrey Scholarship (by St Catherine’s College, Oxford) and Tang Scholarship (by China-Oxford Scholarship Fund). This work is a product of the Global Ecosystems Monitoring (GEM) network (gem.tropicalforests.ox.ac.uk). Fieldwork was funded by grants from the UK Natural Environment Research Council (NE/ 1014705/1) and Advanced Investigator Grants to YM and RV from the European Research Council (GEM-TRAIT and Africa-GHG, respectively). We also acknowledge the Wildlife Division of the Forestry Commission in Ghana as well as the many field assistants who helped with data collection from the field.

## Supplementary data

Supplementary data 1: all_GPP_together_per_site_20221122.csv

Measurements of each carbon cycle component as a mean per plot

Supplementary data 2: all_GPP_together_per_plot_20221122.csv

Measurements of each carbon cycle component as a mean per site

## 10. Supplementary 1 Field measurements and processing procedure

The procedure was written to ensure reproducibility of results and thus includes many processing details.

Leaf area index (LAI) was estimated from hemispherical images taken with a Nikon 5100 camera and Nikon Fisheye Converter FC-E8 0.21x JAPAN near the center of each of the 25 subplots in each plot in each site, at a standard height of 1 m, and during overcast conditions. 22,000 photos were collected in total, every month during 2016-2017(ANK), 2012-2017 (BOB&KOG). Photos were processed using machine learning-based software ‘ilastik’ ^85^ for pixel classification and CANEYE ^86^ for leaf area index calculations. The exposure procedure followed ^87^ and GEM manual ^76^ (http://gem.tropicalforests.ox.ac.uk). The following parameters were supplied to CANEYE.

(1) P1 = angle of view of the fish eye divided by the amount of pixels from centroid of the fish eye circle to where horizon is on the image.
(2) angle of view = 90 degree, in which case, the edge of the photo is the horizon and the centroid of the image is zenith.
(3) COI = 80, consideration of field is 80 degrees, we don’t want the edge of the photo because it is not clear and sometime obscure by tall grasses or saplings.
(4) Sub sample factor =1
(5) Fcover = 20 degree, this is to calculate the percentage of black pixels within central 20-degree ring. We used this to understand the relative openness of canopy for the given image. It is not relevant to LAI
(6) PAIsat = 10, When a pixel is completely black, mathematically, the leaf area index (LAI) is infinite. As we provide CANEYE 25 subplot images for each estimation of LAI, this means all 25 subplot images show black at a given pixel. To address this ‘infinite’issue, we uses a value of 10 for LAI in such cases. This value is based on the guess that, the densest point in a tropical forest should had an LAI of 10.
(7) Latitude 0 and Day of year a random number (not relevant for tropical site LAI)

Then, we extract output from CANEYE using software R. We chose the latest method of LAI calculation offered by CANEYE, ‘CE V6.1 True PAI’. CANEYE reported one LAI value per method (4 methods) per plot per site per month, as a synthesis across 25 subplots images. As systematic error is dominating in LAI calculation, we take the standard deviation of LAI across four methods as the uncertainty for LAI.

Canopy respiration (R_leaf) is calculated as plot-mean LAI multiplied with plot-mean leaf dark respiration (Rdark), a leaf gas exchange measurement. To obtain the leaves, branches for both sun leaves and shade leaves were detached and immediately re-cut under water to restore hydraulic connectivity for subsequent gas exchange measurement. The leaves were fully darkened for 30 min prior to measuring Rdark. Rdark was measured using an open flow gas exchange system (LI-6400XT, Li-Cor Inc., Lincoln, NE, USA) and block temperature was kept constant throughout the sampling period at 30° C. The uncertainty of Rdark was calculated as the standard error of raw measurements ^61^. We convert measurements of Rdark from 30 degree to mean annual air temperature following ^31^. Rdark was measured for sun and shade leaves and from wet to dry seasons. We calculate a basal area community weighted mean for Rdark_sun and Rdark_shade. Then, we calculate canopy respiration per plot using: R_leaf = Rdark_sun * F_sunlit + Rdark_shade* (1 – F_sunlit), where F_sunlit is the sunlit leaf area. It is calculated as Fsunlit = (1 – exp(− K∗LAI))/K where K is the light extinction coefficient ^88^. The final canopy total respiration was calculated as R_leaf * 0.67 to account for daytime light inhibition of leaf dark respiration ^62^.

Above-ground live wood respiration (R_stem), was quantified at monthly intervals by measuring rates of CO2 accumulation to chambers attached to the tree trunk, and scaling using stem surface area allometries, using previous developed equation^89^. Bole respiration per unit surface area was measured using a wood respiration closed dynamic chamber method, from at least 50 trees covering dominating species distributed evenly throughout each plot at 1.3 m height with an IRGA (EGM-4) and soil respiration chamber (SRC-1) connected to a permanent collar. The uncertainty of bole respiration per unit surface area was calculated as the standard error of raw measurements. To recognise the large uncertainty of total stem surface area, mostly due to the simple allometries equation, we assigned an uncertainty of 30%.

Coarse root respiration (R_coarse_root) was not measured, by estimated by R_stem multiplied by 0.21 ± 0.10, following ^82, 90, 91^.

Total soil CO2 efflux (R_soil), called R_soil_no_coarseroot_with_litter in the raw data sheet, was measured every month at the same point in each of the 25 sub-plots on each plot. It was measured using a closed dynamic chamber method with an infra-red gas analyser and soil respiration chamber (EGM-4 IRGA and SRC-1 chamber, PP Systems, Hitchin, UK) sealed to a permanent collar in the soil. Coarse root respiration was assumed missed by the above method ^82^. The uncertainty of R_soil_no_coarseroot_with_litter was calculated as the standard error of raw measurements.

Therefore, the R_soil_no_coarseroot_with_litter is composed of R_rhizosphere (including fine roots, mycorrhizal and exudates) respiration, soil organic matter derived respiration (R_soil_heterotrophic), and soil surface fine litter respiration (R_fine_litter). The percentage of each component was determined by using a partitioning experiment similar to that described in ^84, 92^.

Root exudates NPP was not directly measured while mycorrhizal respiration was incorporated in R_rhizosphere in R_a, bringing uncertainty to NPP and CUE. Following ^84^, We estimated the root exudation rate from literature as (i) 6% of total NPP ^93^ (ii) 59% of root NPP ^94^, (iii) 37% of root respiration (calculated from data in ^95^)

Coarse woody debris respiration and dead wood respiration was not directly measured, which affect the estimates of carbon sink (net ecosystem exchange) but is irrelevant to GPP nor CUE. Study of Amazonia lowland intact forest found CWD respiration as 76% of CWD input, where a steady state (D_cwd = D_cwd_to_soil + R_cwd) was assumed ^96^. However, the proportion of CWD respired could be rather variable ^97^; Recent study at Borneo lowland forest reported a 90% ^46^. In this study, we estimated R_cwd as (0.9+0.76)/2 = 0.83 of D_cwd, with ±0.1 uncertainty.

This study sources stem biomass (or called above ground coarse woody biomass, estimated from tree height and girth) and some NPP components from ^10^. However, it is worth noting that the study is limited in that some minor components (in terms of magnitude) of the carbon cycle were not covered by this study. For instance, Volatile organic compound NPP was found to be very minor components of the carbon cycle of an Amazonian Forest ^62^. Ground flora was neglected in ANK and BOB due to their relatively low abundance, and was included in KOG, a forest to savanna transition zone. Epiphytes and liana were also not counted albeit their wide existence in the field, especially in BOB01.

When combining or multiplying different components of the carbon cycle, uncertainties were propagated following ^62^.

## 11. Supplementary 2: Correlation between NPP and GPP

The following tests were performed as NPP were commonly calculated from GPP with a fixed proportion (known as CUE). The regression shows how well spatial variation of GPP captured the spatial variation of NPP

**Figure S 1.**
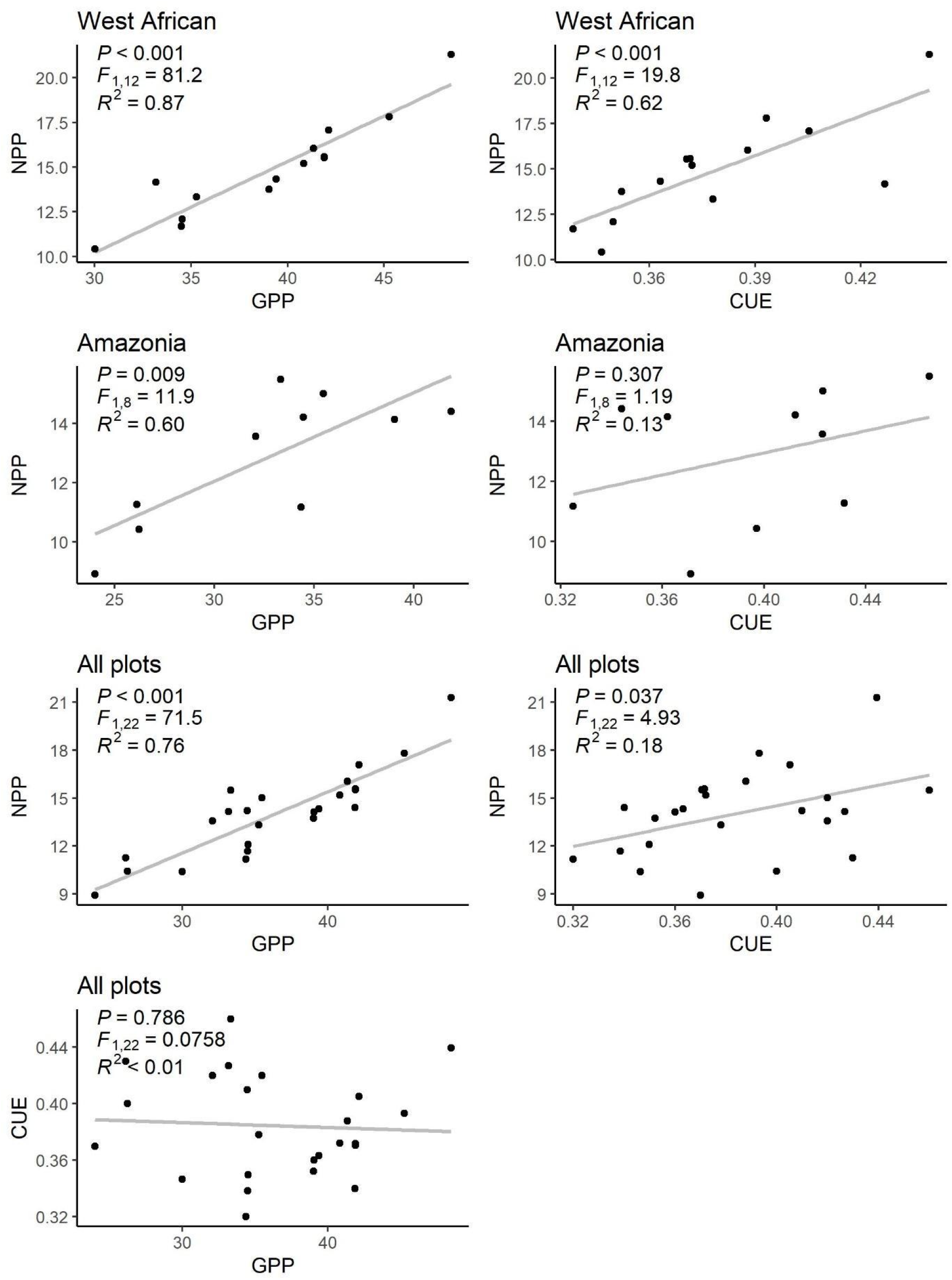
Linear regression between gross primary productivity (GPP) and net primary productivity (NPP) both in unit (Mg C ha-1 yr-1), for West African plots, Amazonia plots and all plots together.

## 12. Supplementary 3 Plots information

**Table S 1.**
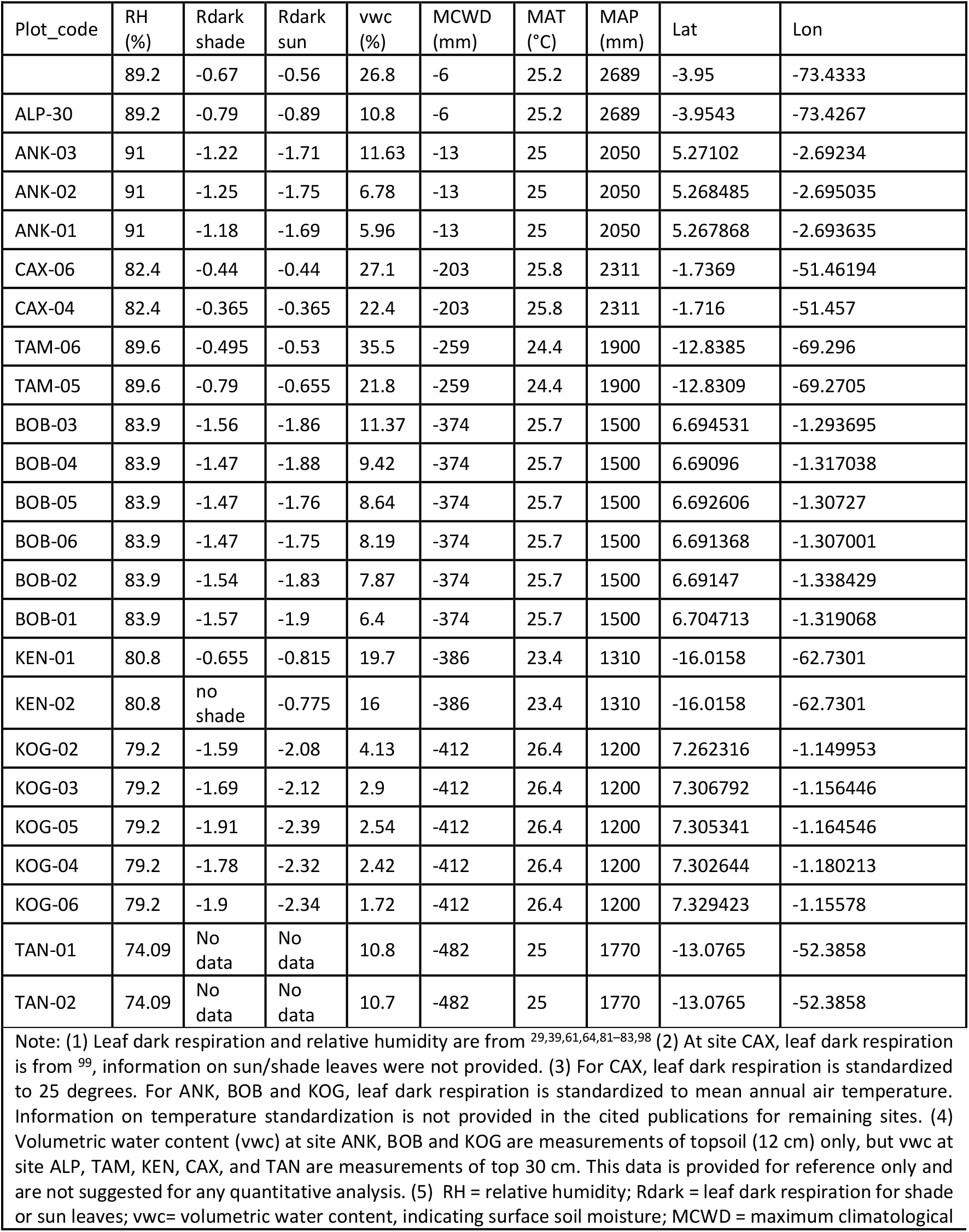
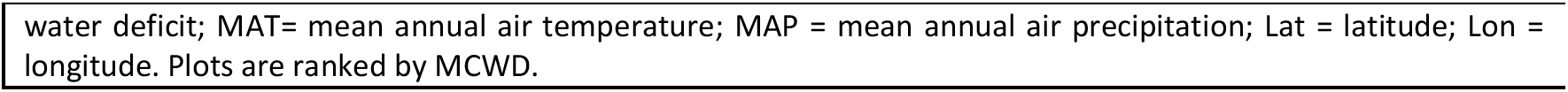
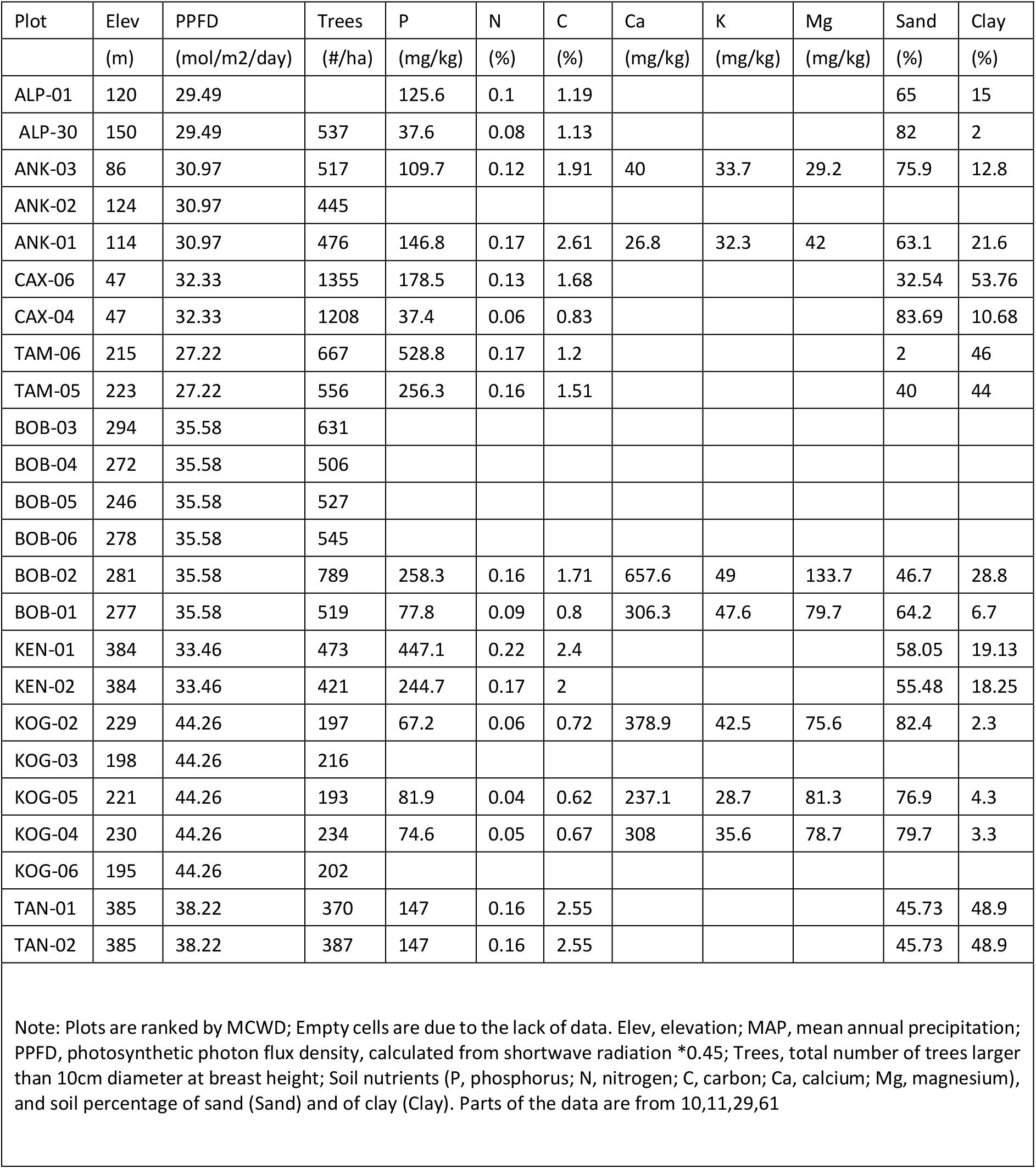
Study plots information. All are one-hectare plots.

## 13. Supplementa ry 4 Thorough carbon budget quantification for West African carbon fluxes

**Figure S 2.**
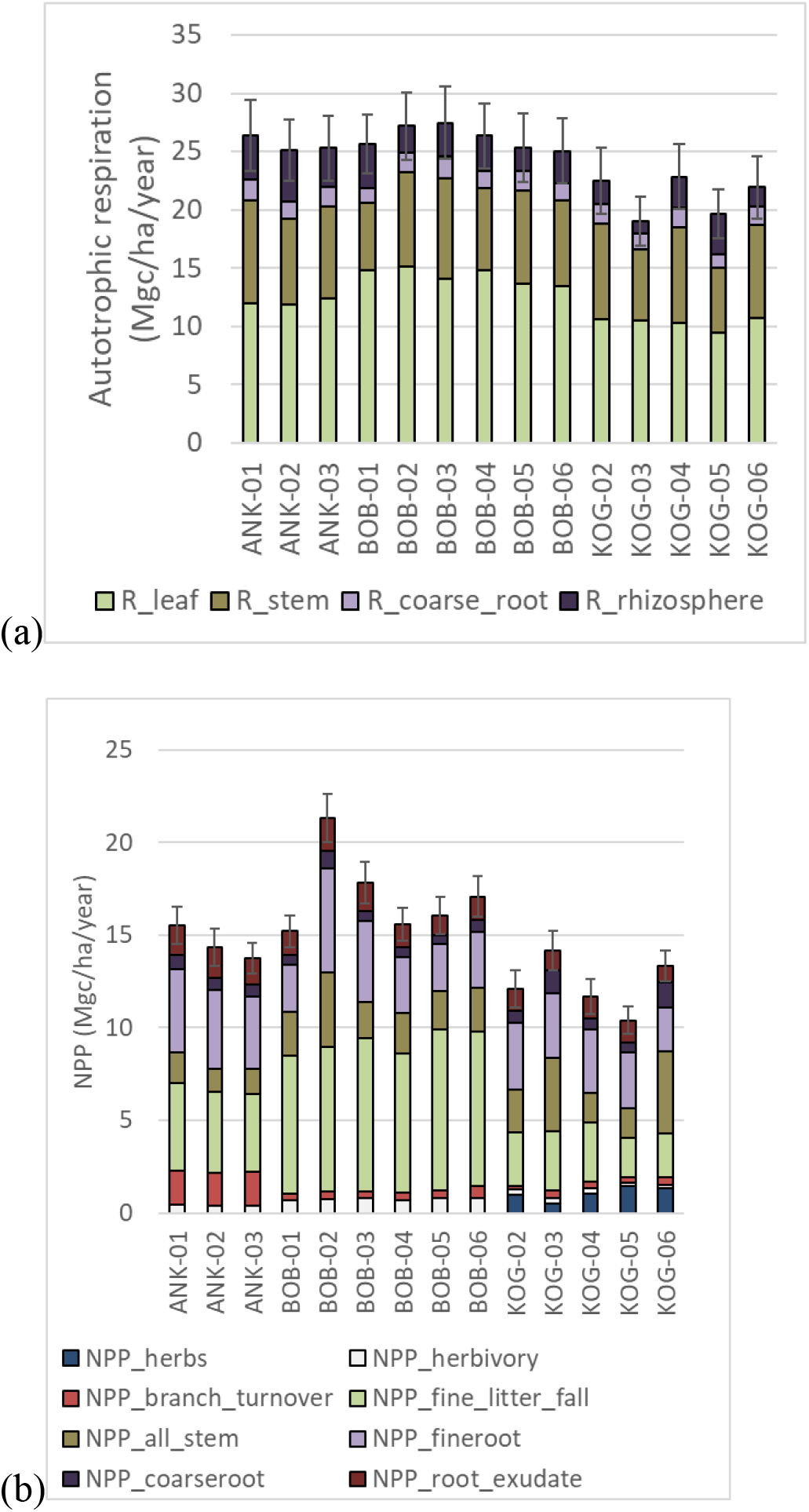
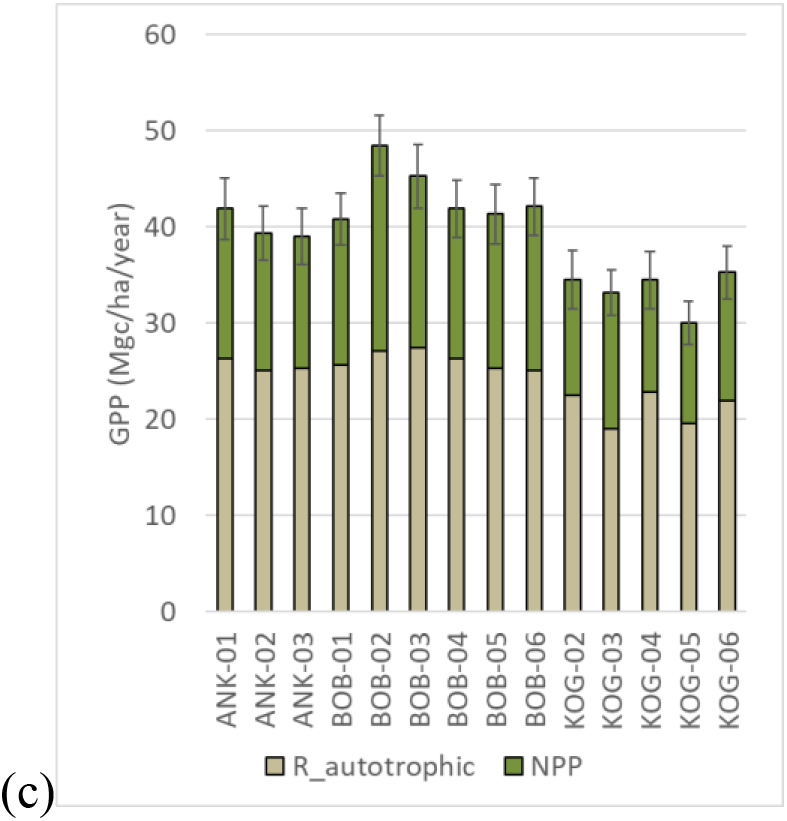
Components of gross primary production (GPP), Autotrophic respiration (R_autotrophic) and net primary production (NPP)

**Figure S 3.**
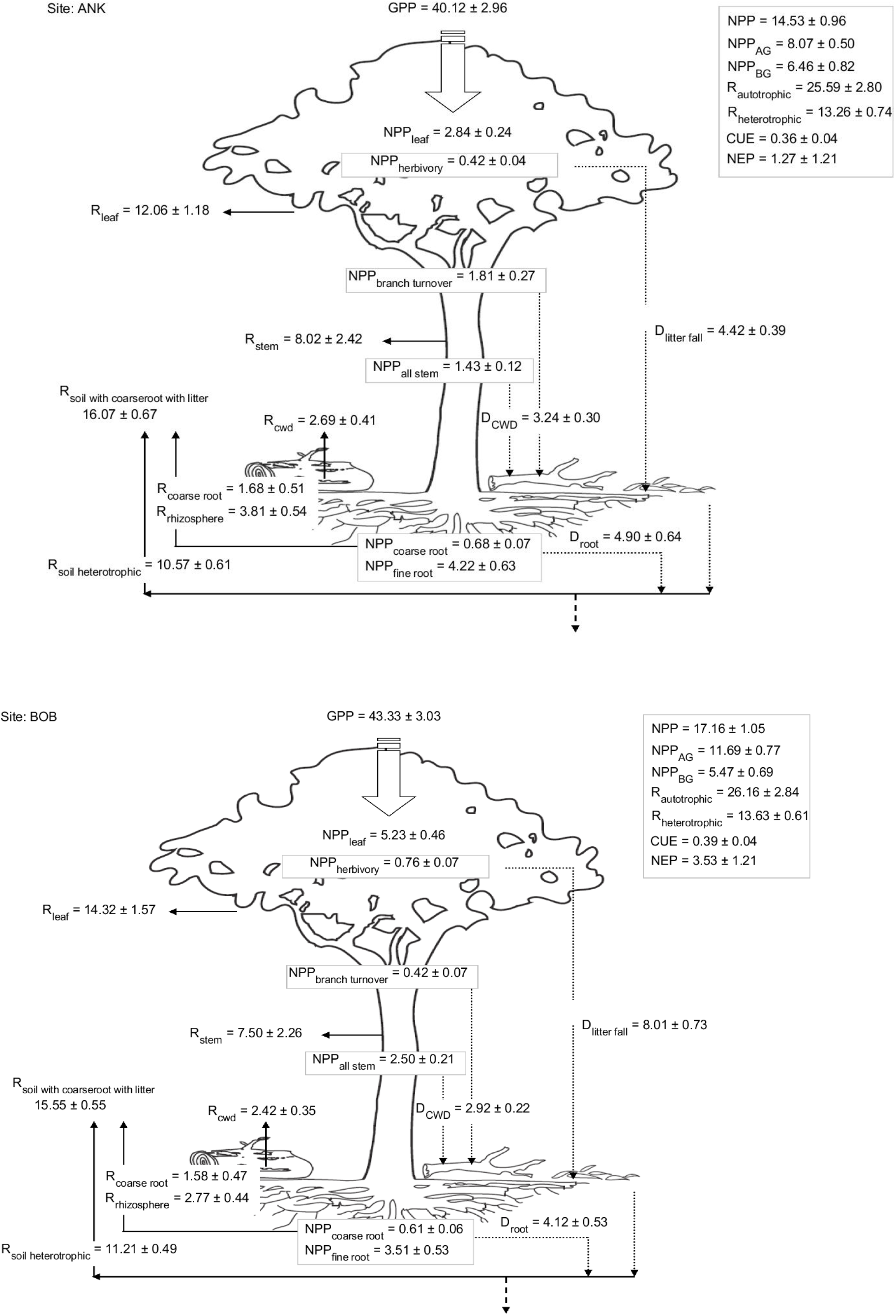
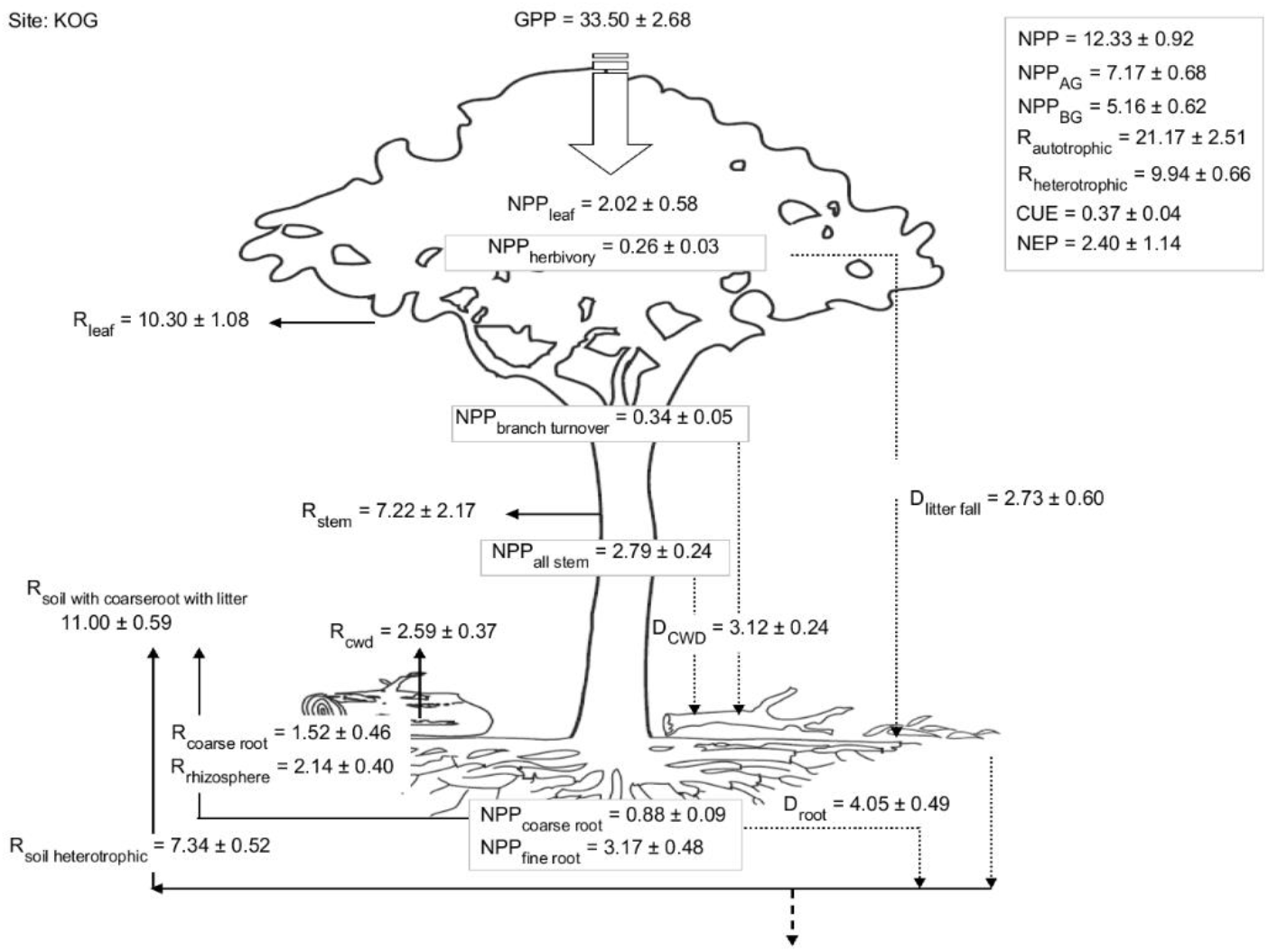
Diagram showing the magnitude and pattern of key carbon fluxes for ANK (3 plots) BOB (6 plots) and KOG (5 plots)

**Table S 2.**
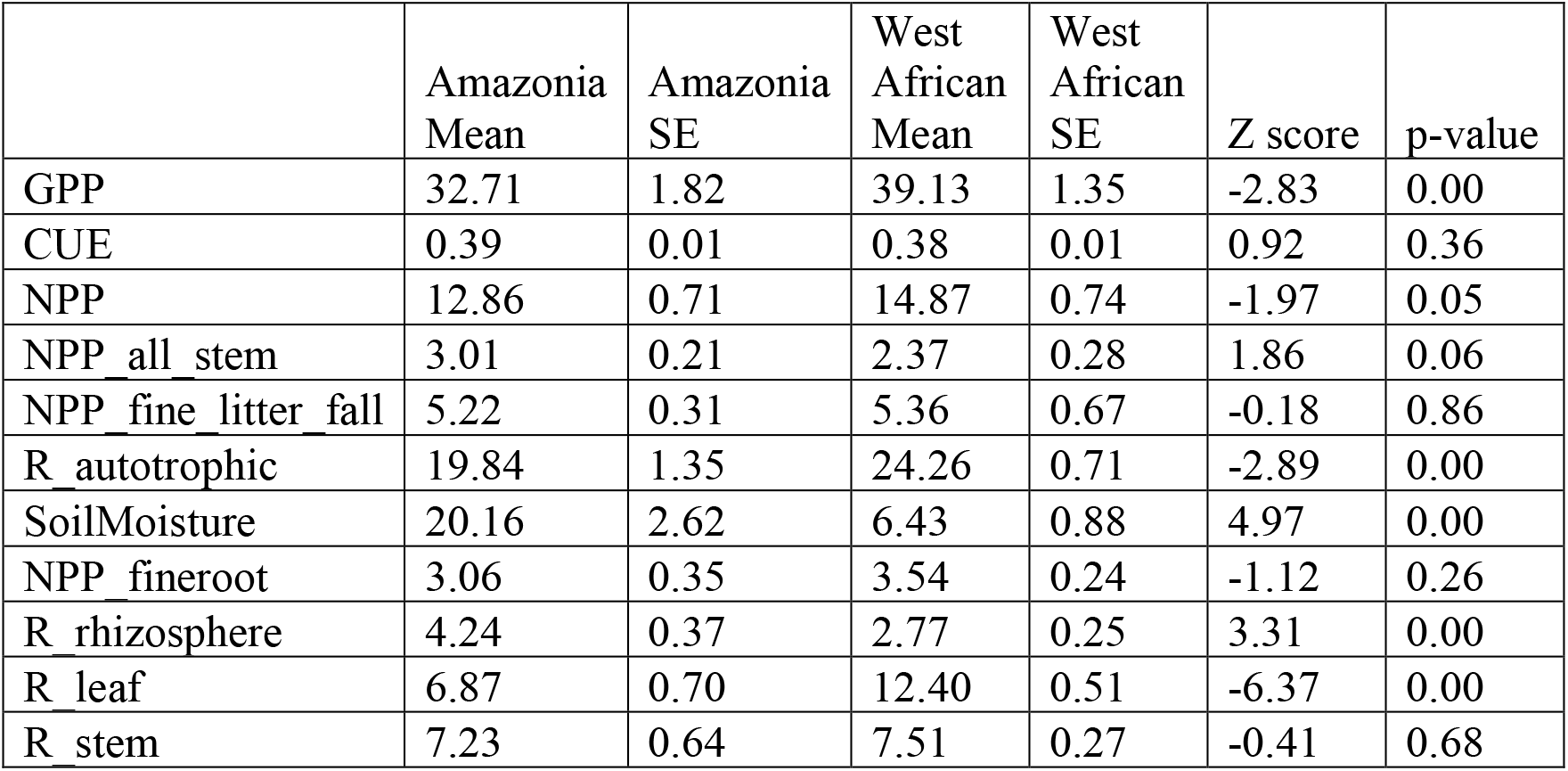
Mean and Standard error (SE) of gross primary production (GPP) and its components across study plots for Amazonia and West African forests, the z-score and p-value test the difference between Amazonia and West African forests. SoilMoisture is soil volumetric water content (%) measured at 12cm depth. See Table 1 for full names and definitions of carbon budget components.

|  | Amazonia<br>Mean | Amazonia<br>SE | West<br>African<br>Mean | West<br>African<br>SE | Z score | p-value |
| --- | --- | --- | --- | --- | --- | --- |
| GPP | 32.71 | 1.82 | 39.13 | 1.35 | -2.83 | 0.00 |
| CUE | 0.39 | 0.01 | 0.38 | 0.01 | 0.92 | 0.36 |
| NPP | 12.86 | 0.71 | 14.87 | 0.74 | -1.97 | 0.05 |
| NPP_all_stem | 3.01 | 0.21 | 2.37 | 0.28 | 1.86 | 0.06 |
| NPP_fine_litter_fall | 5.22 | 0.31 | 5.36 | 0.67 | -0.18 | 0.86 |
| R_autotrophic | 19.84 | 1.35 | 24.26 | 0.71 | -2.89 | 0.00 |
| SoilMoisture | 20.16 | 2.62 | 6.43 | 0.88 | 4.97 | 0.00 |
| NPP_fineroot | 3.06 | 0.35 | 3.54 | 0.24 | -1.12 | 0.26 |
| R_rhizosphere | 4.24 | 0.37 | 2.77 | 0.25 | 3.31 | 0.00 |
| R_leaf | 6.87 | 0.70 | 12.40 | 0.51 | -6.37 | 0.00 |
| R_stem | 7.23 | 0.64 | 7.51 | 0.27 | -0.41 | 0.68 |

**Figure S 4.**
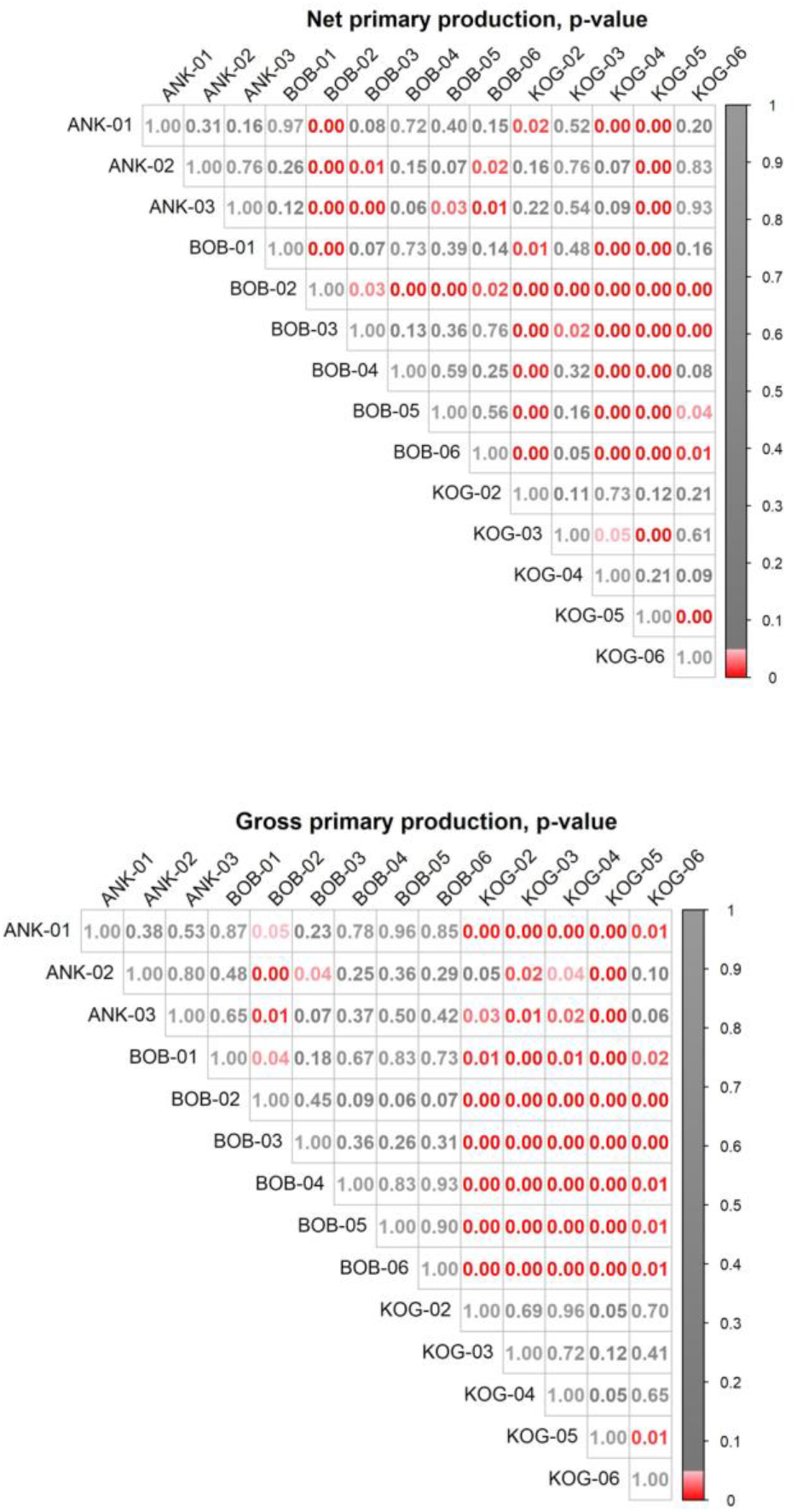
Z-test was used to compare the difference between plots for net primary production (NPP) and gross primary production (GPP). P values between each plot were shown to illustrate significant (red, at 0.05 threshold) and insignificant (grey).

## Notes

### Competing Interest Statement

The authors have declared no competing interest.

